# Structural insights into rice KAI2 receptor provide functional implications for perception and signal transduction

**DOI:** 10.1101/2023.12.28.573577

**Authors:** Angelica M. Guercio, Amelia K. Gilio, Jacob Pawlak, Nitzan Shabek

## Abstract

KAI2 proteins, classified as plant α/β hydrolase receptors, are capable of perceiving smoke-derived butenolide signals and endogenous, yet unidentified KAI2-ligands (KLs). While the number of functional KAI2 receptors varies among land plant species, rice possesses only one KAI2 gene. Rice, a significant crop and representative of grasses, relies on KAI2-mediated Arbuscular mycorrhiza (AM) symbioses to flourish in traditionally arid and nutrient-poor environments. This study presents the first crystal structure of an active rice KAI2 hydrolase receptor. Our structural and biochemical analyses uncover grass-unique pocket residues influencing ligand sensitivity and hydrolytic activity. Through structure-guided analysis, we identify a specific residue whose mutation enables the increase or decrease of ligand perception, catalytic activity, and signal transduction. Furthermore, we investigate rice KAI2-mediated signaling by examining its ability to form a complex with its binding partner, the F-box protein DWARF3 (D3) ubiquitin ligase, demonstrating the significant role of hydrophobic interactions in the KAI2-D3 interface. This study provides new insights into the diverse and pivotal roles of the KAI2 signaling pathway in the plant kingdom, particularly in grasses.

## Introduction

Rice (*Oryza sativa*) is an important crop and is often considered a representative for other grasses, which include many additional staple crops like maize (*Zea mays*), wheat (*Triticum aestivum*), and sorghum (*Sorghum bicolor*). Symbiosis with Arbuscular mycorrhiza (AM) fungi is crucial for many plants across all lineages, but its importance in grasses is especially relevant since grasses are often grown in nutrient-poor soils and AM can assist with the uptake of nitrogen and phosphate (1). Additionally, because grasses are often found in ecosystems prone to drought stress, AM symbiosis can aid in drought tolerance by improving water-use efficiency (2, 3). AM fungi also increase tolerance to biotic and abiotic factors like disease (4) and salinity (5). The more tolerant and acclimatory grasses can be, the larger yields they can produce, critical for agriculture because of their value and importance as staple crops.

In 2015, it was discovered amongst mounting knowledge of fungi-derived signal perception in plants (6), the receptor protein D14-Like (D14L, often termed KAI2 or HTL in other species, referred to as KAI2 hereafter) is necessary for rice AM colonization (7). Additionally, KAI2 is part of a protein signaling pathway that was discovered to sense external signaling molecules from the soil called karrikins (KARs) (8). KARs are molecules produced by the combustion of plant material, that are able to descend into the soil in post-fire rain and subsequently act as bioactive germination stimulants (9, 10). KARs were shown to stimulate germination in 1200 species of plants across 80 genera, including non-fire relevant plants like rice (11, 12). Identification via reverse genetics screens in Arabidopsis and rice of the genes involved in KAR signaling revealed three currently known proteins: the receptor KAI2 (KARRIKIN INSENSITIVE 2), an E3 ubiquitin ligase F-box protein D3/MAX2 (DWARF3/MORE AXILLARY GROWTH2), and a substrate that is ubiquitinated and degraded as a result of the pathway, SMAX1/SMXL2 (SUPPRESSOR OF MAX2-1 and SMAX1-LIKE 2) (8, 13, 14). These three proteins share homology and identity with proteins in the signaling pathway of the plant hormone strigolactone (SL) (15–20). SL is a phytohormone found in plants whose perception signaling pathway as currently known relies on a homologous receptor to KAI2, D14 (DWARF14), the same D3/MAX2 E3 ligase, and a homologous substrate SMAX1-LIKE 6, 7, and 8 (SMXL) in Arabidopsis or D53 (DWARF53) in rice (15–20). In fact, based on evolutionary analyses, KAR signaling genes were found to be ancestral to SL genes (21, 22). In addition to the evolutionary context of KAR and SL signaling pathways, there is mounting evidence that the KAR signaling pathway exists to sense a yet unknown endogenous small molecule thusly called KL (KAI2 ligand) (23, 24). This includes the striking phenotypes of *kai2*, *max2*, and/or *smax1* mutants even in the absence of KAR. There is not only crosstalk between the KAR and SL signaling pathways in terms of their significance in plants, but specifically as both of these signaling pathways influence the ability of AM to colonize plant roots.

SL signaling modulates many physiological traits in plants including shoot branching, leaf growth, leaf senescence, secondary stem thickening, formation of adventitious roots, lateral roots, and root hairs (25–28). The list of SL-dependent phenotypes continues to grow as their roles are studied in diverse species and contexts (29). The process by which SL and AM symbioses overlap occurs first when SLs are exuded by plant roots (30). These SLs are perceived by the fungi in the soil by an unknown mechanism; in response, SLs activate fungal metabolism and hyphae branching. (31, 32). However, the KAR signaling pathway plays an even greater role; perception and symbioses with AM fungi requires the KAR signaling pathway. Loss of function *d14l/kai2* in rice abolishes any symbiotic activity (33). The same is true in loss of function rice mutants of the E3 ligase *d3* which targets SMAX1 (34). It was found from *smax1* loss of function rice that SMAX1 is a negative regulator of AM symbioses. As such *smax1* loss of function rice increases colonization and suppresses the *kai2* and *d3* mutant phenotypes (35). The *smax1* loss of function also induces expression of AM- response genes and increases SL exudation and biosynthesis (35).

The specific signaling molecule perceived by the KAI2 protein to initiate symbioses is unknown, but by studying the receptor itself we can accelerate the search for a fungal ligand. The KAI2/D14 family of receptors has been studied with the aid of structural biology in numerous species. in In *Striga hermonthica* structural studies have provided insights into the most sensitive of these family of receptors and their responses towards a wide variety of ligands (36). Structural studies in Arabidopsis and pea have given insights into ligand-binding, hydrolysis dynamics, and the residues necessary for coordinating small molecules in the pocket (37, 38). Mutating pocket residues has been shown to be an effective method to alter ligand sensitivity in vitro and *in vivo* (37–40) aiding in the exploration of receptor function and evolution/co-evolution with KLs.

Ex vivo complex formation of KAI2s with the E3 ligase D3 is challenging outside of yeast 2 hybrid systems (41–43). However, in SL signaling the complex of D14 with D3 has been determined (44, 45). Previous work examining the conservation of D14/KAI2 family residues along the receptor-D3 interface highlighted several residues conserved across the entire family that could contribute to that interface but have not yet been explored (46).

Here we present the novel structure of the KAI2 receptor in rice and use this structure to inform mutational optimization of the receptor towards synthetic ligands. We also employ rice KAI2 as a model to better explore its interface with the E3 ligase D3. This study will provide broader insights into the mechanism by which KAI2 perceives ligands and induces the formation of symbiotic relationships with fungi as well as emphasize the value of rice KAI2 as a representative for other important grasses.

## Results

### The structure of rice KAI2 reveals a conserved ligand-binding pocket

To obtain greater elucidation of the mode of ligand perception, we determined the crystal structure of rice (*Oryza sativa*, Os) KAI2 to 1.28 Å resolution (Fig. 1 and Table S1). As observed in previously determined KAI2 crystal structures, OsKAI2 comprises of a canonical α/β-hydrolase fold with the binding pocket situated within the central core domain, beneath the helical lid domain (residues Tyr125-Ser197) (Fig. 1A and Fig. S1). A Dali structural similarity search revealed that OsKAI2 has greatest similarity to Arabidopsis (At) KAI2 with a sequence identity of 77% and RMSD of 0.5 Å over 268 Cα (Fig. S2).

**Figure 1:**
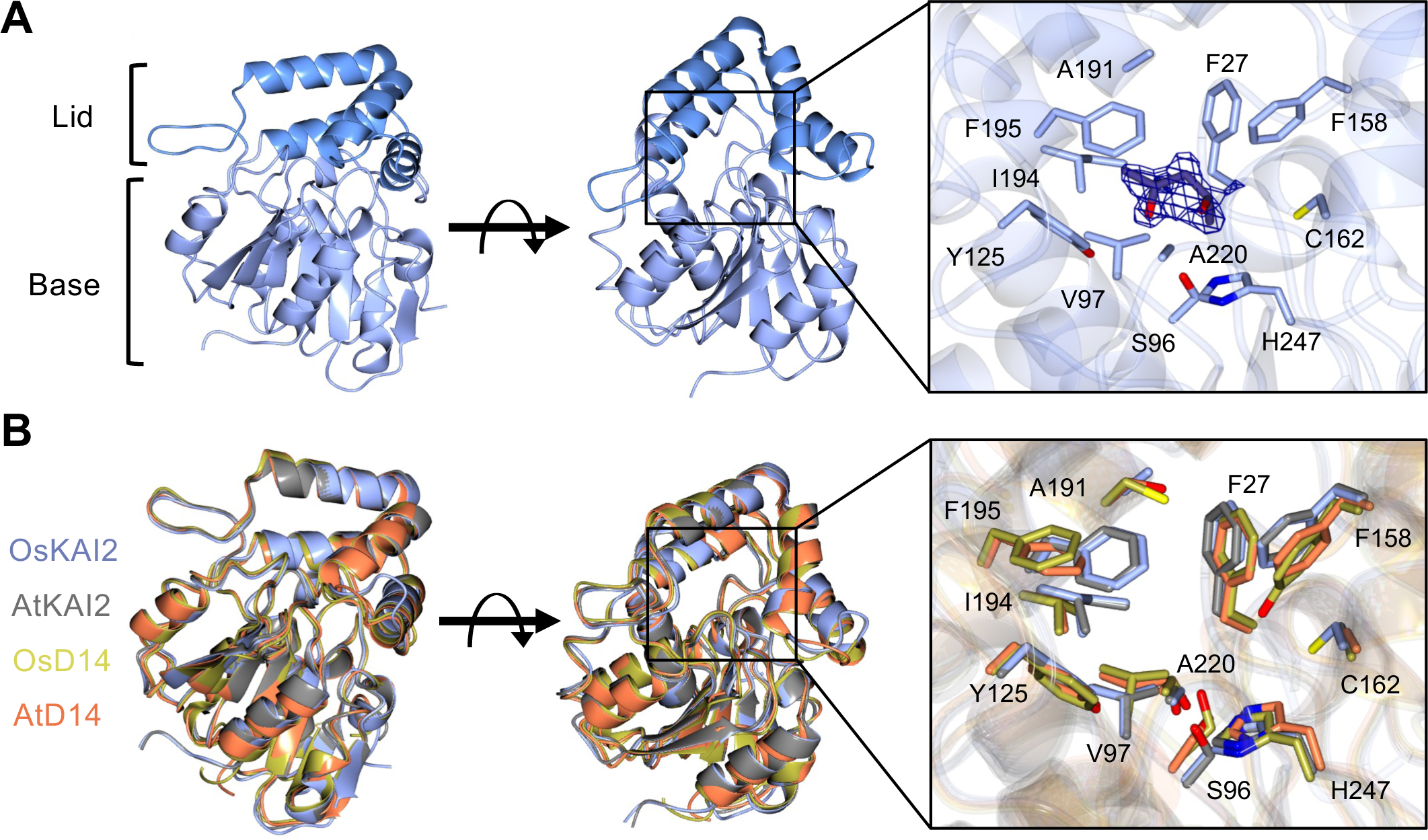
Overview of rice KAI2 crystal structure and comparison with other relevant α/β hydrolases. **(A)** Overall structure of rice (Os) KAI2 with MPD bound in the active site. Lid and Base domains can be observed from the side view with lid shown in dark blue and base in light blue. The top view of the structure can be observed upon rotation. MPD is bound in the active site with all surrounding residues labeled, showing the catalytic S96 and H247 at the base of the active site and the pocket residue of interest, C162 to the right. The 2Fo-Fc map of the density surrounding MPD is shown to sigma =1.0. **(B)** Superposition of OsKAI2 (blue) with AtKAI2 (grey), OsD14 (gold) and AtD14 (orange) illustrating high similarity. Both global and pocket superpositions are illustrated.

Upon a closer inspection of the binding pocket, the conserved Ser-His-Asp catalytic triad is positioned at the base of the active site. The pocket comprises of a hydrophobic cleft made up of several phenylalanine, isoleucine, and valine residues, making it well suited to bind the aromatic rings of KAR or KL-type ligands. Interestingly, we have also determined the OsKAI2 crystal structure with 2-Methyl-2,4-pentanediol (MPD), originating from the crystallization reagents (Table S1). MPD is bound within the pocket, forming hydrogen bonding interactions with the key catalytic residues H247 and S96, alongside Y125. While MPD is not the natural ligand, its orientation within the pocket may indicate how the endogenous ligand could interact with the pocket residues.

The OsKAI2 pocket was compared against that of AtKAI2 which shows similar pocket residues, with the exceptions of C162 and A191 in OsKAI2, that represent smaller side-chain groups in the positions of A161 and G191 in AtKAI2 (Fig. 1B and Fig. S1). The OsKAI2 and AtKAI2 structures were also compared to the analogous receptors for SL signaling in both rice OsD14, and in the model plant Arabidopsis, AtD14 (Fig. 1). To that end, we determine a new AtD14 structure at a higher resolution (2 Å) than the previously reported structures and was hence applied here for comparative analysis (Fig. 1, Table S1). Structural superposition with OsKAI2 showed an RMSD of 1.2Å over 265 Cα and 1.2Å over 264 Cα for OsD14 and AtD14 respectively, and both D14s gave a sequence identity of 56%, confirming high structural similarity in all four proteins examined here. When focusing on the binding pocket, several residues including the Ser-His-Asp catalytic triad are conserved in all proteins, highlighting its importance in the catalytic mode of action performed by α/β hydrolases. However, there are some clear differences in both the positioning and type of residue found in the pocket when comparing between the KAI2s and D14s. Most noticeably, F27/F26 in OsKAI2 and AtKAI2 are shifted by 1.2 Å in respect to the positioning of F28 in both D14s (Fig. 1 and Fig. S1). Similarly, F195 in both KAI2s is shifted into the pocket by 2.7 Å when compared to F195 in both D14s. Furthermore, both KAI2s possess Y125/124 in place of F126 in D14s, with the hydroxyl group reaching 1.3 Å further into the pocket; similarly, both KAI2s possess the larger hydrophobic I194 in place of V194 seen in the D14s. Conversely, we found that A220 in both rice and Arabidopsis KAI2s is replaced with S220 in D14s which is slightly larger more polar, perhaps indicating a role in strigolactone binding (Fig. 1B and Fig. S1). The OsKAI2 pocket analyses revealed a pocket measured to have a volume of 108 Å (47). This is an intermediate pocket size when compared to others in the family, such as AtKAI2 with an approximate volume of 63 Å, OsD14 at 149 Å, and ShHTL7 at 215 Å (Fig. S2). Collectively, the differences between KAI2 and D14s demonstrate that many residues reaching further into the pocket in the KAI2s, suggesting an adapted and more compact pocket to better fit KL/SL like ligands.

### Evolutionary context of rice KAI2 indicates highly conserved grass receptors

In order to understand how KAI2 in rice has evolved within the context of the KAI2 family, a maximum likelihood tree was generated from an amino acid alignment of 131 KAI2s and one D14 (Fig. 2 and Table S2). Noticeably, based on the tree topology, KAI2s within grasses are very similar to one another. Consequently, the gene encoded DNA sequences were examined for increased selection; using the RELAX method comparing the grasses branches to the rest of the tree, the test for selection intensification (K = 30.08) was significant (p = 0.000, LR = 22.00) (48). As such, it is clear that grasses are experiencing selective pressures, hence we employed structural studies to take a deeper look at grass-unique residues as compared to the rest of the tree. Residues within the ligand binding pocket, as illustrated in Fig. 1 and Fig. S1, encompass the invariant catalytic triad S96 and H247. While residues F27, V97, Y125, F158, I194, F195, and A220 show some variability in other species, they exhibit minimal variations and maintain high conservation across the entire family, without specific divergence in grasses. Residue A191 has been previously reported to be a divergent residue in other species but is an alanine at that position in the majority of species assessed including in grasses. Previous studies have demonstrated how amino acid substitutions in the ligand binding pocket can influence receptor function by altering specificity and efficacy towards different ligands (37, 40, 49, 50). Strikingly, due to its distinct proximity to the ligand binding pocket, residue C162 was selected for further inspection (Fig. 1 and Fig. 2). We investigated the amino acid identities at position 162 in all species assessed within the phylogenetic tree. The pie charts plotted illustrate the proportion of each amino acid in grasses (Fig. 2, top right), the divergent family of receptors in striga (top left), and in all other representative species (Fig. 2, bottom left). Notably, upon examining all sequences, alanine emerges as the most common amino acid at position 162 (or its equivalent when aligned). This consistency is evident not only in Arabidopsis KAI2 but also in 73% of all species examined outside of grasses and striga (Fig. 2 and Fig. S1). Intriguingly, when examining some of the most active KAI2 receptors in *Striga hermonthica*, a distinct set of residues at this position has diverged, comprising leucine, valine, isoleucine, and methionine (Fig. 2). The receptor HTL7, which demonstrates heightened sensitivity to various tested ligands both in vivo and in vitro, features leucine in this position (Fig. S1). We next examined how these residues affect the activity of rice KAI2 as a receptor and enzyme by generating the substitutions C162A and C162L.

**Figure 2.**
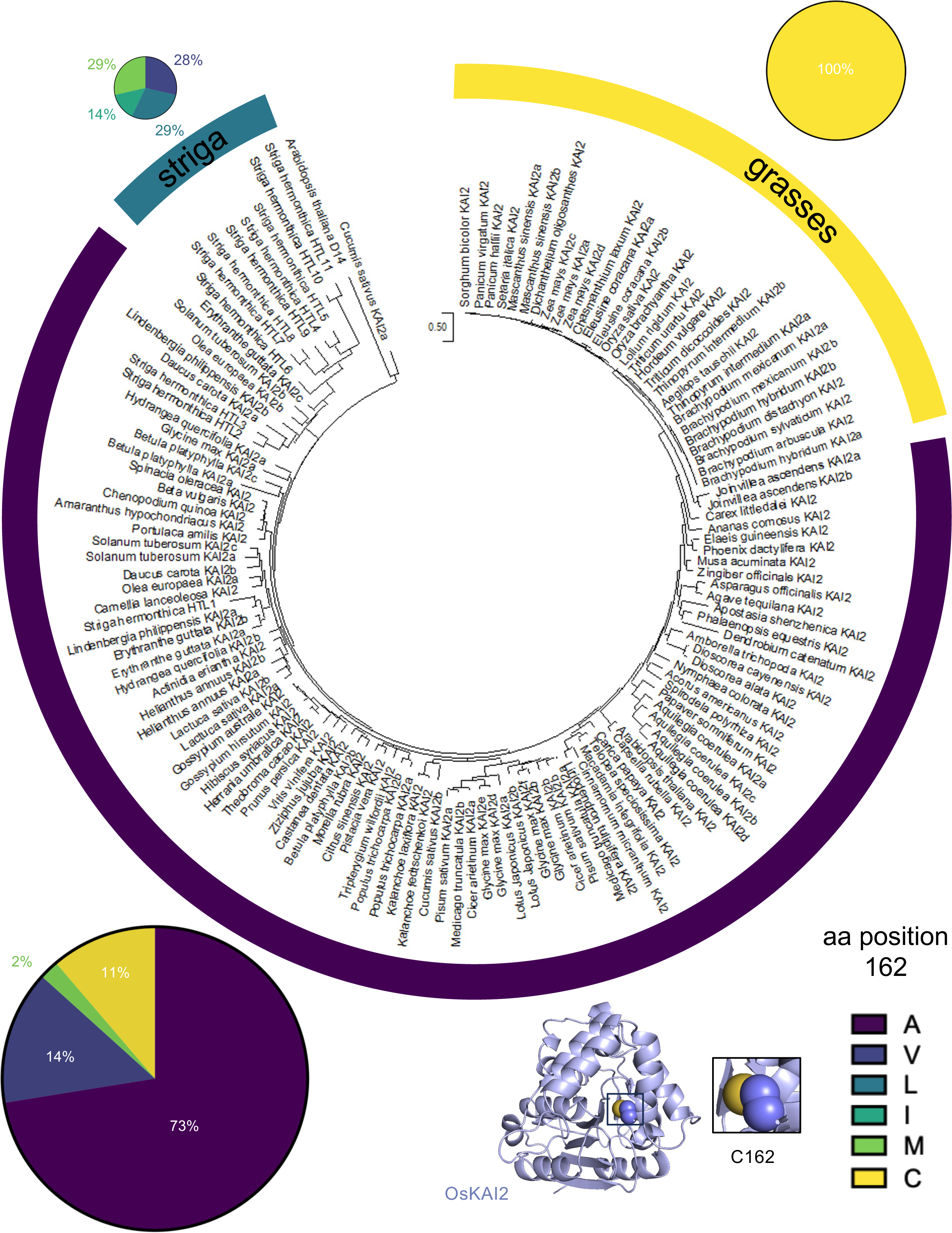
Evolutionary context of rice KAI2 and analyses of site 162. Maximum likelihood tree of 132 amino acid sequences. Pie charts are split to encompass three groups highlighted on tree including grasses (top right), striga (top left), and all other species (bottom left). Pie charts represent the proportion of species within that group with amino acids A, V, L, I, M, or C respectively at site 162. Site 162 is detailed on OsKAI2 structure (bottom middle). Area of pie chart is proportional to the number of species sampled.

### Perturbing the rice KAI2 binding pocket affects sensitivity towards synthetic ligands

In the absence of knowledge of the endogenous KL, synthetic ligands have proven immensely effective in better understanding the KAI2 family receptors. Specifically, KAI2 receptors have been observed to be extremely enantio-specific (37, 51). As such, we tested the response of the wildtype OsKAI2 and the C162A and C162L mutants (Fig. 3A, D, and G, Fig. S3A-C) towards both the (+)-GR24 and (-)-GR24 enantiomers (synthetic SL analogs). Generally, SL receptor D14 responds to (+)-GR24 due to its similarity to natural SLs, whilst KAI2 proteins tend to respond to (-)-GR24 more readily (37, 40, 52, 53). Indeed, using differential scanning fluorimetry (DSF), we observed no response with any KAI2 examined to the (+)-GR24 (Fig. S3D-F). Markedly, the wildtype (WT) OsKAI2 responds to increasing concentrations of (-)-GR24 as shown by a shift in the melting temperature (Tm) curve (Fig. 3B). Interestingly, in comparison to the WT OsKAI2, the C162A mutation seems to slightly stabilize the protein as seen by the relative rightward shift in Tm (52° C compared to 47° C); nonetheless, the mutation to alanine led to a loss of all sensitivity towards (-)-GR24 (Fig. 3E). This is likely due to the increased rigidity of OsKAI2-C162A, which compromises ligand-induced destabilization normally observed in these receptors. Furthermore, the C162L mutation provides an increased sensitivity towards (-)-GR24 as illustrated by at least a 2-fold increase in sensitivity (62.5 vs 125 μM) when compared to the WT KAI2 (Fig. 3B and H).

**Figure 3.**
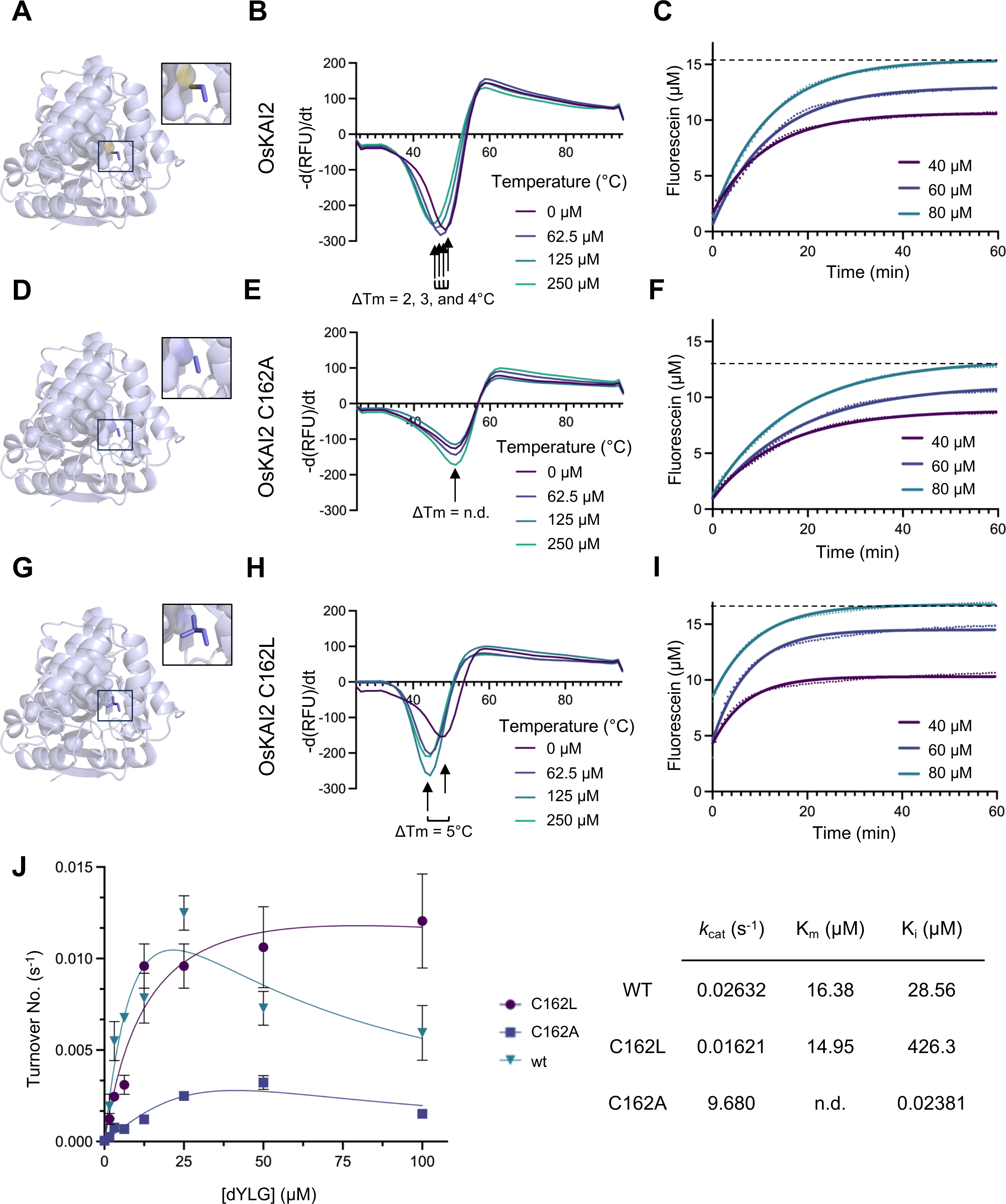
Mutational analyses of rice KAI2 and its effect on receptor/enzymatic activity. Wildtype KAI2 **(A-C)**, KAI2 C162A **(D-F)**, and KAI2 C162L **(G-H)** were subjected to differential scanning fluorimetry **(B, E, H)** and dYLG hydrolysis assays **(C, F, I)**. Structures are shown as cartoons with residue at position 162 shown as sphere **(A, D, G)**. dYLG hydrolysis **(C, F, I)** colored lines represent non-linear regression curved fit, based on average values of triplicates of raw data (shown in dots). Dashed lines represent plateau value for highest concentration of dYLG (80µM). dYLG hydrolysis initial rate kinetics assay **(J)** was used to determine Kcat, Km, and Ki values in table.

We next investigated the impact of the pocket mutations on receptor function by measuring enzyme activity towards pro-fluorescent synthetic ligands. To that end, we employed hydrolysis assay using the fluorogenic derivative of Yoshimulactone Green (YLG (54)) which lacks a methyl group on the lactone ring (desmethyl YLG, dYLG (55)). As expected, WT OsKAI2 is able to effectively hydrolyze dYLG with a typical exponential to plateau curve as previously seen in this family of enzymes (Fig. 3C). OsKAI2-C162A mutant is less effective at hydrolyzing the ligand than WT as evidenced by the shallower hydrolysis curves and lower plateau (Fig. 3F). Strikingly, the C162L mutant surpasses the wildtype in activity, reaching a plateau at a faster rate than observed with the wildtype (Fig. 3I). Kinetic analysis of the initial rates of hydrolysis for the two mutants and WT OsKAI2 further corroborate these observations (Fig. 3J). In fact, we observe substrate inhibition for both the WT and C162A mutant as the enzyme turnover numbers drop at high ligand concentrations. However, C162L shows negligible inhibition and reaches the greatest turnover numbers, suggesting that a leucine at this position can enhance the enzyme activity. This observation has been previously hypothesized by the residue presence in a range of the most active receptors in this family.

### Insights into rice KAI2-D3 interface

The KAR/KL pathway relies on the ubiquitin proteasome system to trigger transcriptional repressor degradation upon ligand perception through the recruitment by the F-box protein D3 as part of the SCF ubiquitin ligase module). As such, we wanted to explore the interaction between OsKAI2 and OsD3. To that end, we generated a 3D model of the complex using the homologous SL complex AtD14 with OsD3-ASK1 as was previously reported (46, 56). By modeling the OsKAI2 into the complex, we were able to identify four distinct highly conserved amino acids that are involved in KAI2/D14 and D3 interaction: D30, D51, E172, and N181 (Fig. 4A and B). These residues are within a 2.7-3.5 Å distance to potential interactors across the protein interface and hence could participate in forming hydrogen bonds and salt bridges between the two proteins (Fig. 4B). Subsequently, we mutated these four polar residues into small hydrophobic alanine to observe the effects on protein binding (OsKAI2^int^, Fig. S4). We explored the change in the binding surface’s hydrophobicity upon mutating the four interface residues to alanine and found that indeed the hydrophobicity of the surface was substantially increased (Fig. 4C). Interestingly, upon interrogation by pull-down assays, we consistently found that the OsKAI2^int^ mutant bound OsD3-ASK1 better than WT (Fig. 4D). This indicates that hydrophobic interactions can increase the receptor-E3 complex (KAI2-D3) interaction, which may not be highly favorable under physiological conditions, where plasticity and dissociation are required for successful signal transduction.

**Figure 4.**
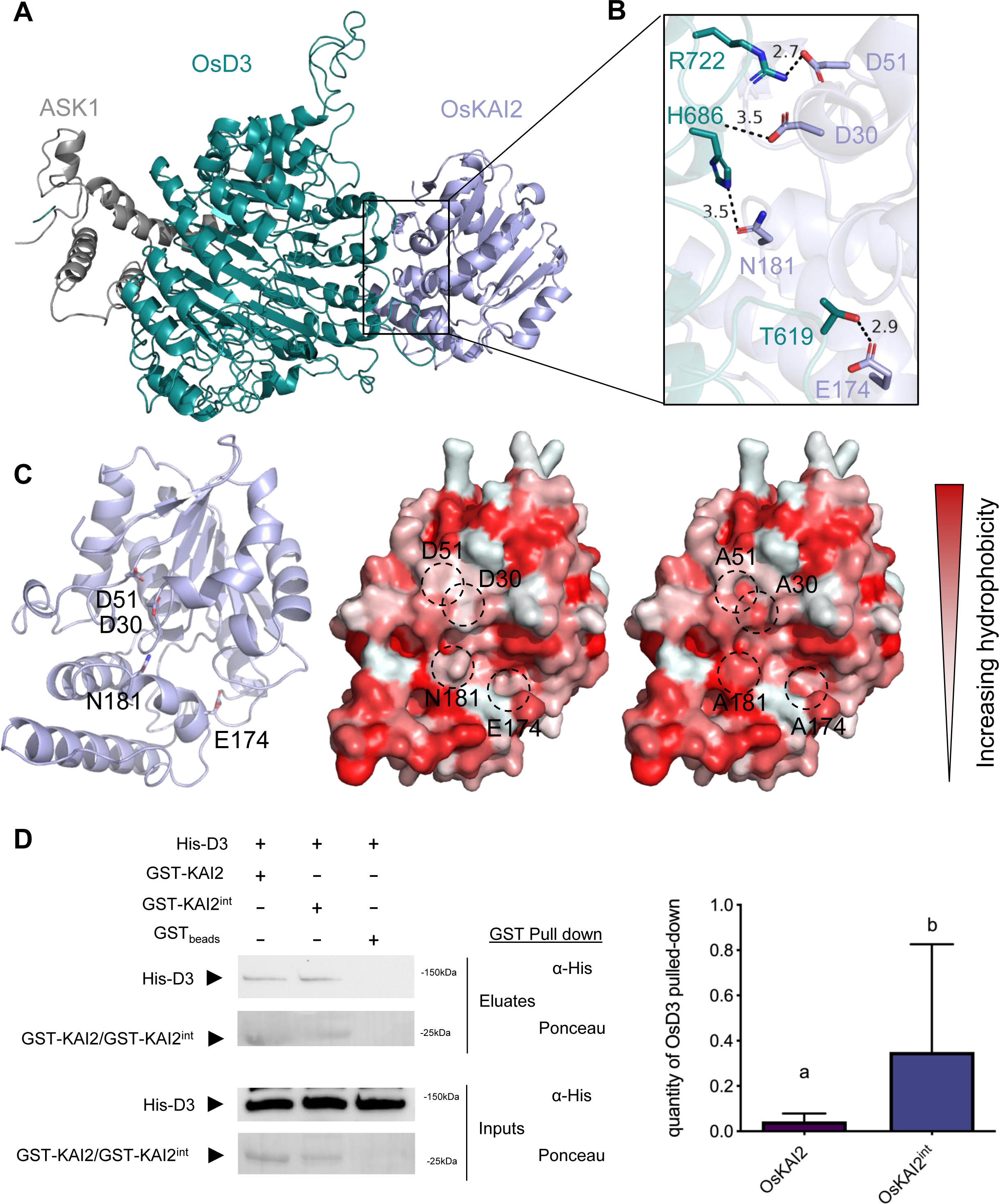
KAI2-MAX2 interaction analysis. **(A)** 3D model of OsD3-ASK1 and OsKAI2 complex (model was calculated based on PDB 5ZHG). **(B)** Zoom in view of polar residues interacting across the interface. Angstrom distances between side chains are shown in dash lines and were calculated by PyMOL. **(C)** Left: molecular architecture of OsKAI2 determined in this study shown in cartoon (light blue) and the interface residues are shown and labeled (black). Middle and right: surface representation of OsKAI2 WT structure (middle) and KAI2 mutant (KAI2^int^, right) with hydrophobicity levels shown in shades of red. Dashed circles indicate the D3-KAI2 interface polar residues that are mutated to alanine (D51A, D30A, N181A, E174A). **(D)** Left: Pull-down experiment with GST-KAI2 or GST-KAI2^int^ mutant with His-MSB tagged OsD3-ASK1. Right: quantification of D3 pull-down by GST-KAI2 or GST-KAI2^int^ mutant over three independent experiments.

## Discussion

KAI2 receptors play important regulatory roles in numerous physiological processes. A more comprehensive understanding of the structure of KAI2 receptors offers deeper insights into their function in perceiving and processing both exogenous and endogenous signals, and the consequent impact on symbioses and plant development. Rice, a significant crop and representative of grasses, depends on KAI2-mediated Arbuscular mycorrhiza (AM) symbioses to thrive in traditionally arid and nutrient-poor environments. In this study, we determined the first crystal structure of an active rice KAI2 hydrolase receptor and explore its function. The importance of several residues in the ligand-binding pocket have been of great interest including the residue 160 (position 161 in rice), which is positioned adjacent to the residue 162 examined here, and was shown to be important in differentiating ligand sensitivity between lotus and pea duplicated receptors (37, 40). Other studies looking across all asterid KAI2s also identified the 161 residue (position 162 in rice, examined here) as one of the four residues defining the two broad classes of KAI2s (Y124 and F124-types) (39). These four sites (positions 96, 124, 139, and 161) were previously tested for their function in Arabidopsis swapping between Y124 and F124 residues as single, double, triple, and quadruple mutants. The V161A mutation (which aligns with the C162A and C162L mutations in this study) alone was enough to significantly decrease hypocotyl elongation inhibition, but substitutions to other amino acids at this site were not examined thus far. Here, we examined the position 162 in response to (-)-GR24 and dYLG ligands. Interestingly, an alanine at this site nearly removes all sensitivity, while a cysteine is sensitive to both, and a leucine is the most sensitive towards both ligands tested. Taken together, our data not only corroborate previous reports but also reveal certain critical positions within the binding pocket of grasses KAI2s that evolved to sense and probably select highly specific yet unidentified KL.

The downstream regulation of KAI2 mediated signaling has been examined here through the interaction with its known binding partner, MAX2/D3 F-box protein. Through structure-guided analysis, we have generated KAI2-D3 model with high probability based on D14-D3 crystal structure. Mutating pivotal polar residues within the KAI2-D3 interface significantly amplifies the binding between the proteins, indicating that increased hydrophobicity generates a more tightly bound and rigid complex. While this data elucidates the precise interface between KAI2-D3, it also suggests that naturally occurring polar residues within this interface may contribute to less rigidity and weaker interactions, characteristic of E3 ligases and their binding partners.

Future studies, both in planta and at the structure-function levels, along with the identification of KL in grasses, will contribute to elucidating the precise network of interacting residues with the E3 ligase, D3, as well as with other binding partners, such as the SMAX1 transcriptional co-repressor.

## Materials and Methods

### Protein preparation and purification

AtKAI2, OsKAI2, the C162A mutant and the C162L mutant were independently cloned and expressed as 6× His-SUMO fusion proteins from the expression vector pAL (Addgene). BL21 (DE3) cells transformed with the expression plasmid were grown in LB broth at 16 °C to an OD600 of _∼_0.8 and induced with 0.25 mM IPTG for 16 h. Cells were harvested, re-suspended and lysed in extract buffer (50 mM Tris, pH 8.0, 200 mM NaCl, 5 mM imidazole, 4% Glycerol). All His-SUMO-proteins were isolated from soluble cell lysate by Ni-NTA resin. The proteins were eluted with 250 mM imidazole and subjected to anion-exchange. The eluted proteins were then cleaved with TEV (tobacco etch virus) protease overnight at 4 °C. The cleaved His-SUMO tag was removed by passing through Ni-NTA resin. All proteins were concentrated by ultrafiltration to 3–10 mg/mL−1.

### Crystallization, data collection and structure determination

The crystals of OsKAI2 were grown at 25 °C by the hanging-drop vapor diffusion method with 1.0 μL purified protein sample (at 12mg/mL) mixed with an equal volume of reservoir solution containing 0.1M Sodium HEPES, 0.1M MOPS (acid) pH 7.5, 0.018M Magnesium chloride hexahydrate, 0.018M Calcium chloride dihydrate, 9.4% v/v MPD, 9.4% v/v PEG 1000, 9.4% v/v PEG 3350. The crystals of AtD14 were grown at 25 °C by the hanging-drop vapor diffusion method with 1.0 μL purified protein sample mixed with an equal volume of reservoir solution containing 0.15M Ammonium acetate, 0.01M Calcium chloride dihydrate, 0.1M Tris pH 8.5, 28% broad PEG smear from BCS (Molecular Dimensions), 20% NPD. Crystals of maximum size were obtained and harvested from the reservoir solution. X-ray diffraction data was integrated and scaled with HKL2000 package (57). OsKAI2 crystal structures were determined by molecular replacement using the AtKAI2 model (PDB: 5Z9H) (58) as the search model. AtD14 crystal structure was determined by molecular replacement using the PDB 41H4 model. All structural models were manually built, refined, and rebuilt with PHENIX (59) and COOT (60).

### Alignments and Phylogenetic Analyses

Amino acid sequences were retrieved from NCBI, Phytozome, and GenBank with accession information listed in Table S1. Multiple sequence alignment was performed in MEGA X version 10.2.6 (61) using the ClustalW algorithm (62) with a final alignment consisting of 393 positions. To identify best fitting model for phylogeny construction, MEGA X model testing was performed and the amino acid substitution model with the lowest BIC score (Bayesian Information Criterion) chosen for analyses was the LG+G model with 5 discrete gamma categories. The evolutionary history was inferred by using the Maximum Likelihood method and LG matrix-based model with 5 discrete gamma categories. The evolutionary history was inferred by using the Maximum Likelihood method and Le_Gascuel_2008 model (63). The tree with the highest log likelihood (−16612.94) is shown. The percentage of trees in which the associated taxa clustered together is shown next to the branches. Initial tree(s) for the heuristic search were obtained automatically by applying Neighbor-Join and BioNJ algorithms to a matrix of pairwise distances estimated using the JTT model, and then selecting the topology with superior log likelihood value. A discrete Gamma distribution was used to model evolutionary rate differences among sites (5 categories (+G, parameter = 0.6363). This analysis involved 132 amino acid sequences. There were a total of 393 positions in the final dataset. For selection analyses RELAX (48) was used in the Datamonkey (64) server with coding sequences for 100 sequences listed in Table S1. 32 sequences from evolutionary analyses were removed because of their extensive gapping. 342 sites were examined for codon dn/ds among the grass branches compared to the whole tree and showed intensification of selection compared to the whole tree.

### Thermal Shift Assays

Differential Scanning Fluorimetry (DSF) experiments were performed on a CFX96 Touch^TM^ Real-Time PCR Detection System (Bio-Rad Laboratories, Inc., Hercules, California, USA) using excitation and emission wavelengths of 560-590 and 610-650 nm, respectively. SYPRO® Orange Protein Gel Stain (Sigma Aldrich) was used as the reporter dye. Samples were heat-denatured using a linear 25 to 95 °C gradient at a rate of 1.3 °C per minute after incubation at 25 °C for 30 min in the absence of light. The denaturation curve was obtained using CFX manager™ software. Final reaction mixtures were prepared in triplicate in 96-well white microplates, and each reaction was carried out in 30 μL scale in reaction buffer (20mM HEPES pH 7.3, 250mM NaCl, 1mM TCEP) containing 20uM protein, 0-250 μM ligand (as shown on the Fig. 3B, E, and H), 2% (v/v) DMSO, and 10x Sypro Orange (from 5000x manufacturer’s stock). Plates were incubated in darkness for 30 min before analysis. In the control reaction, DMSO was added instead of ligand. All experiments were repeated three times.

### dYLG hydrolysis

dYLG (desmethyl Yoshimulactone Green; gifted from Dr. Mark Waters, UWA) hydrolysis assays were conducted in reaction buffer (50LmM MES pHL6.0, 150 mM NaCl, and 1 mM DTT) in a 50 μl volume on a 96-well, F-bottom, black plate (Greiner Bio-One, Monroe, NC, USA). The final concentration of dimethyl sulfoxide (DMSO) was equilibrated for all samples to final concentration of 0.8%. The intensity of the fluorescence was measured by a Synergy|H1 Microplate Reader (Agilent Technologies, Santa Clara, CA, USA) with excitation by 480 nm and detection by 520 nm. Readings were collected using 13-s intervals over 60 min. Background auto-YLG hydrolysis correction was performed for all samples. Raw fluorescence data were converted directly to fluorescein concentration using a standard curve. Data that were generated in Excel were transferred to GraphPad Prism 9 for graphical analysis. One-way ANOVA was performed to compare each condition with a post hoc Tukey multiple comparison test. All experiments were run with technical triplicates, and independent experiments were performed three times.

### Initial rates kinetic assays

dYLG activity assays were conducted in 50 μl of reaction buffer (50 mM MES pH 6.0, 250 mM NaCl and 1 mM DTT) in a 96-well, F-bottom, black plate (Greiner), in accordance with the manufacturer’s instructions. 2uM protein was added to each well containing 0-80 µM dYLG and shaken for 5 s before data was collected. The intensity of the fluorescence was measured using a Synergy|H1 Microplate Reader (Agilent Technologies, Santa Clara, CA, USA) with excitation at 480 nm and detection at 520 nm. Readings were collected at 2 s intervals over 2 min. The rate of YLG hydrolysis was determined from the initial linear portion of this fluorescence data in Excel and were transferred to Prism Graphpad v.10 for kinetic analysis. Kinetic constants were determined through non-linear regression based on substrate inhibition kinetics using Prism Graphpad v.10.

### Pulldown assay

OsKAI2 and the OsKAI2^int^ mutant were independently cloned and expressed as GST (Glutathione-S-Transferase) fusion protein from the expression vector pCOOL (Addgene). BL21 (DE3) cells transformed with the expression plasmid were grown in LB broth at 16°C to an OD600 of ∼1.0 and induced with 0.2mM IPTG for 16 h. Cells were harvested, re-suspended and lysed in extract buffer (50mM Tris, pH 8.5, 200 mM NaCl, 5mM DTT, 8% glycerol). Following cell lysis, clarification, and centrifugation, the cell lysate was incubated with GST beads for 1 h at 4°C. Beads were then washed twice with wash buffer containing 50 mM Tris, 150 mM NaCl, 1% Glycerol and 1 mM TCEP. Beads were then washed once with 0.025% BSA blocking solution and twice with wash buffer. For control, GST beads were incubated with 0.05% BSA. Protein-bound GST beads were incubated with purified His-MSB OsD3-ASK1 and 100 μM GR24ent5DS (or Acetone as control) for 30 min on ice. Following two washes with wash buffer, elution was achieved with 50 mM Tris, 10 mM Reduced glutathione pHL7.2, and 5 mM DTT. After addition of fourfold concentrated sample buffer, boiled samples were resolved via SDS-PAGE, and proteins were visualized using Ponceau stain and Western blot with monoclonal anti-His (Invitrogen MA1-21315), and polyclonal anti-GST (Thermo Fisher Scientific, Waltham, MA, USA, CAB4169) antibodies. Quantification of pulled-down OsD3 was measured by band intensity compared to the ladder 150kDa molecular weight marker for 3 independent experiments. T-test was performed to compare pull-down for wildtype compared to KAI2^int^ mutant.

### Structural modelling and structure visualization

Sequence for OsKAI2 was threaded through existing AtD14–OsD3-ASK1 complex structures using SWISS-MODEL with PDB ID: 5HZG as template. Residues of importance were identified as being within hydrogen bonding distance of potential hydrogen bond donor/acceptor pairs between OsKAI2 and OsD3. The 3D structure illustration and analysis were generated using PyMOL Molecular Graphics System, Schrödinger, LLC. Hydrophobicity was assessed using hydrophobicity scale as defined by (65).

## Acknowledgements

N.S. is supported by the National Science Foundation (NSF-CAREER Award #2047396, NSF-EAGER Award #2028283, and Award #2139805), and by the U.S. Department of Energy, Office of Science, Biological and Environmental Research, Genomic Science Program grant no. DE-SC0023158. We thank the beamline staff at the Advanced Light Source (U.S. DOE Office of Science User Facility under Contract No. DE-AC02-05CH11231, is supported in part by the ALS-ENABLE program funded by the National Institutes of Health, National Institute of General Medical Sciences, grant P30 GM124169-01).

## Author Contributions

A.M.G., A.K.G and N.S. conceived and designed the experiments. A.M.G., A.K.G and J.P. conducted the protein purification and biochemical assays. A.M.G conducted crystallization experiments. Structural, and functional, analyses were performed by A.M.G and A.K.G. A.M.G wrote the manuscript with help from A.K.G, J.P. and N.S.

## Disclosure Statement

N.S. has an equity interest in Oerth Bio and serves on the company’s Scientific Advisory Board.

The work and data submitted here have no competing interests, or other interests that might be perceived to influence the results and/or discussion reported in this paper.

## Data and materials availability

The atomic coordinates of structures of OsKAI2, OsKAI2 with MPD and AtKAI2 were deposited in the Protein Data Bank with accession codes 8VCZ, 8VD1, 8VD3. All relevant data are available from the corresponding author upon request.

## Notes

### Competing Interest Statement

The authors have declared no competing interest.

### Summary of Updates

Author list is updated on the online server

## References

1. Smith, S. E., Jakobsen, I., Grønlund, M., and Smith, F. A. (2011) Roles of arbuscular mycorrhizas in plant phosphorus nutrition: interactions between pathways of phosphorus uptake in arbuscular mycorrhizal roots have important implications for understanding and manipulating plant phosphorus acquisition Plant physiology 156, 1050–1057,

2. Abdalla, M., Bitterlich, M., Jansa, J., Püschel, D., and Ahmed, M. A. (2023) The role of arbuscular mycorrhizal symbiosis in improving plant water status under drought Journal of Experimental Botany 74, 4808–4824 10.1093/jxb/erad249

3. Li, J., Meng, B., Chai, H., Yang, X., Song, W., Li, S. et al. (2019) Arbuscular mycorrhizal fungi alleviate drought stress in C3 (Leymus chinensis) and C4 (Hemarthria altissima) grasses via altering antioxidant enzyme activities and photosynthesis Frontiers in Plant Science 10, 499,

4. Campos-Soriano, L., García-Martínez, J., and San Segundo, B. (2012) The arbuscular mycorrhizal symbiosis promotes the systemic induction of regulatory defence-related genes in rice leaves and confers resistance to pathogen infection Mol Plant Pathol 13, 579–592 10.1111/j.1364-3703.2011.00773.x

5. Alqarawi, A. A., Abd Allah, E., and Hashem, A. (2014) Alleviation of salt-induced adverse impact via mycorrhizal fungi in Ephedra aphylla Forssk Journal of Plant Interactions 9, 802–810,

6. Gutjahr, C., Banba, M., Croset, V., An, K., Miyao, A., An, G. et al. (2008) Arbuscular mycorrhiza-specific signaling in rice transcends the common symbiosis signaling pathway Plant Cell 20, 2989–3005 10.1105/tpc.108.062414

7. Gutjahr, C. (2015) Rice perception of symbiotic arbuscular mycorrhizal fungi requires the karrikin receptor complex Science 350, 10.1126/science.aac9715

8. Waters, M. T. (2012) Specialisation within the DWARF14 protein family confers distinct responses to karrikins and strigolactones in Arabidopsis Development 139, 10.1242/dev.074567

9. Flematti, G. R., Ghisalberti, E. L., Dixon, K. W., and Trengove, R. D. (2004) A compound from smoke that promotes seed germination Science 305, 977–977 10.1126/science.1099944

10. Van Staden, J., Jäger, A. K., Light, M. E., and Burger, B. V. (2004) Isolation of the major germination cue from plant-derived smoke South African Journal of Botany 70, 10.1016/S0254-6299(15)30206-4

11. Dixon, K. W., Merritt, D. J., Flematti, G. R., and Ghisalberti, E. L. (2009) Karrikinolide - A phytoreactive compound derived from smoke with applications in horticulture, ecological restoration and agriculture Acta Horticulturae 813,

12. Kulkarni, M., Sparg, S., Light, M., and Van Staden, J. (2006) Stimulation of rice (Oryza sativa L.) seedling vigour by smoke_-_water and butenolide Journal of agronomy and crop science 192, 395–398,

13. Nelson, D. C. (2011) F-box protein MAX2 has dual roles in karrikin and strigolactone signaling in Arabidopsis thaliana Proc Natl Acad Sci 108, 10.1073/pnas.1100987108

14. Stanga, J. P., Smith, S. M., Briggs, W. R., and Nelson, D. C. (2013) SUPPRESSOR OF MORE AXILLARY GROWTH2 1 Controls Seed Germination and Seedling Development in Arabidopsis Plant Physiology 163, 318–330 10.1104/pp.113.221259

15. Arite, T., Umehara, M., Ishikawa, S., Hanada, A., Maekawa, M., Yamaguchi, S., and Kyozuka, J. (2009) D14, a strigolactone-Insensitive mutant of rice, shows an accelerated outgrowth of tillers Plant and Cell Physiology 50, 1416–1424-1416–1424 10.1093/pcp/pcp091

16. Jiang, L., Liu, X., Xiong, G., Liu, H., Chen, F., Wang, L. et al. (2013) DWARF 53 acts as a repressor of strigolactone signalling in rice Nature 504, 401–405-401–405 10.1038/nature12870

17. Soundappan, I., Bennett, T., Morffy, N., Liang, Y., Stanga, J. P., Abbas, A. et al. (2015) SMAX1-LIKE/D53 Family Members Enable Distinct MAX2-Dependent Responses to Strigolactones and Karrikins in Arabidopsis The Plant Cell 27, 3143–3159 10.1105/tpc.15.00562

18. Zhou, F., Lin, Q., Zhu, L., Ren, Y., Zhou, K., Shabek, N. et al. (2013) D14-SCF D3 - dependent degradation of D53 regulates strigolactone signalling Nature 504, 406–410 10.1038/nature12878

19. Gao, Z., Qian, Q., Liu, X., Yan, M., Feng, Q., Dong, G. et al. (2009) Dwarf 88, a novel putative esterase gene affecting architecture of rice plant Plant Molecular Biology 71, 265–276,

20. Liu, W., Wu, C., Fu, Y., Hu, G., Si, H., Zhu, L. et al. (2009) Identification and characterization of HTD2: a novel gene negatively regulating tiller bud outgrowth in rice Planta 230, 649–658,

21. Bythell-Douglas, R. (2017) Evolution of strigolactone receptors by gradual neo-functionalization of KAI2 paralogues BMC Biol 15, 10.1186/s12915-017-0397-z

22. Delaux, P. M., Xie, X., Timme, R. E., Puech-Pages, V., Dunand, C., Lecompte, E., et al. (2012) Origin of strigolactones in the green lineage New Phytologist 10.1111/j.1469-8137.2012.04209.x

23. Conn, C. E. (2015) Convergent evolution of strigolactone perception enabled host detection in parasitic plants Science 349, 10.1126/science.aab1140

24. Conn, C. E., and Nelson, D. C. (2016) Evidence that KARRIKIN-INSENSITIVE2 (KAI2) Receptors may Perceive an Unknown Signal that is not Karrikin or Strigolactone Frontiers in Plant Science 6, 1–7 10.3389/fpls.2015.01219

25. Brewer, P. B., Koltai, H., and Beveridge, C. A. (2013) Diverse roles of strigolactones in plant development In Molecular Plant,

26. Smith, S. M., and Li, J. (2014) Signalling and responses to strigolactones and karrikins Current Opinion in Plant Biology 21, 23–29 10.1016/j.pbi.2014.06.003

27. Ruyter-Spira, C., Al-Babili, S., van der Krol, S., and Bouwmeester, H. (2013) The biology of strigolactones In Trends in Plant Science,

28. Bennett, T., and Leyser, O. (2014) Strigolactone signalling: Standing on the shoulders of DWARFs Current Opinion in Plant Biology 22, 7–13 10.1016/j.pbi.2014.08.001

29. Samynathan, R., Venkidasamy, B., Shariati, M. A., Muthuramalingam, P., and Thiruvengadam, M. (2023) Biosynthesis, functional perspectives, and agricultural applications of strigolactones Brazilian Journal of Botany 1-20,

30. Cook, C. E., Whichard, L. P., Turner, B., Wall, M. E., and Egley, G. H. (1966) Germination of witchweed (striga lutea lour.): Isolation and properties of a potent stimulant Science 154, 1189–1190 10.1126/science.154.3753.1189

31. Besserer, A., Puech-Pagès, V., Kiefer, P., Gomez-Roldan, V., Jauneau, A., Roy, S., et al. (2006) Strigolactones stimulate arbuscular mycorrhizal fungi by activating mitochondria PLoS biology 4, e226,

32. Akiyama, K., Matsuzaki, K., and Hayashi, H. (2005) Plant sesquiterpenes induce hyphal branching in arbuscular mycorrhizal fungi NATURE 435, 824–827 10.1038/nature03608

33. Gutjahr, C., Gobbato, E., Choi, J., Riemann, M., Johnston, M. G., Summers, W. et al. (2015) Rice perception of symbiotic arbuscular mycorrhizal fungi requires the karrikin receptor complex Science 350, 1521–1524 10.1126/science.aac9715

34. Yoshida, S., Kameoka, H., Tempo, M., Akiyama, K., Umehara, M., Yamaguchi, S. et al. (2012) The D3 F_-_box protein is a key component in host strigolactone responses essential for arbuscular mycorrhizal symbiosis New Phytologist 196, 1208–1216,

35. Choi, J., Lee, T., Cho, J., Servante, E. K., Pucker, B., Summers, W., et al. (2020) The negative regulator SMAX1 controls mycorrhizal symbiosis and strigolactone biosynthesis in rice Nature Communications 11, 10.1038/s41467-020-16021-1

36. Toh, S. (2015) Structure-function analysis identifies highly sensitive strigolactone receptors in Striga Science 350, 10.1126/science.aac9476

37. Guercio, A. M., Torabi, S., Cornu, D., Dalmais, M., Bendahmane, A., Le Signor, C. et al. (2022) Structural and functional analyses explain Pea KAI2 receptor diversity and reveal stereoselective catalysis during signal perception Communications Biology 5, 10.1038/s42003-022-03085-6

38. Yao, J. (2018) An allelic series at the KARRIKIN INSENSITIVE 2 locus of Arabidopsis thaliana decouples ligand hydrolysis and receptor degradation from downstream signalling Plant J 96, 10.1111/tpj.14017

39. Martinez, S. E., Conn, C. E., Guercio, A. M., Sepulveda, C., Fiscus, C. J., Koenig, D. et al. (2022) A KARRIKIN INSENSITIVE2 paralog in lettuce mediates highly sensitive germination responses to karrikinolide Plant Physiology 1–17 10.1093/plphys/kiac328

40. Carbonnel, S., Torabi, S., Griesmann, M., Bleek, E., Tang, Y., Buchka, S. et al. (2020) Lotus japonicus karrikin receptors display divergent ligand-binding specificities and organ-dependent redundancy PLoS Genetics 16, 10.1371/journal.pgen.1009249

41. Toh, S., Holbrook-Smith, D., Stokes, M. E., Tsuchiya, Y., and McCourt, P. (2014) Detection of parasitic plant suicide germination compounds using a high-throughput Arabidopsis HTL/KAI2 strigolactone perception system Chemistry & biology 21, 988–998 10.1016/j.chembiol.2014.07.005

42. Holbrook-Smith, D., Toh, S., Tsuchiya, Y., and McCourt, P. (2016) Small-molecule antagonists of germination of the parasitic plant Striga hermonthica Nature Chemical Biology 12, 10.1038/nchembio.2129

43. Khosla, A., Morffy, N., Li, Q., Faure, L., Chang, S. H., Yao, J. et al. (2020) Structure–Function analysis of SMAX1 reveals domains that mediate its karrikin-induced proteolysis and interaction with the receptor KAI2 Plant Cell 32, 2639–2659 10.1105/tpc.19.00752

44. Yao, R. (2016) DWARF14 is a non-canonical hormone receptor for strigolactone Nature 536, 10.1038/nature19073

45. Shabek, N., Ticchiarelli, F., Mao, H., Hinds, T. R., Leyser, O., and Zheng, N. (2018) Structural plasticity of D3–D14 ubiquitin ligase in strigolactone signalling Nature 563, 652–656 10.1038/s41586-018-0743-5

46. Tal, L., Guercio, A. M., Varshney, K., Young, A., Gutjahr, C., and Shabek, N. (2023) C-terminal conformational changes in SCF-D3/MAX2 ubiquitin ligase are required for KAI2-mediated signaling New Phytol 239, 2067–2075 10.1111/nph.19101

47. Tian, W., Chen, C., Lei, X., Zhao, J., and Liang, J. (2018) CASTp 3.0: computed atlas of surface topography of proteins Nucleic Acids Res 46, W363–w367 10.1093/nar/gky473

48. Wertheim, J. O., Murrell, B., Smith, M. D., Kosakovsky Pond, S. L., and Scheffler, K. (2014) RELAX: Detecting Relaxed Selection in a Phylogenetic Framework Molecular Biology and Evolution 32, 820–832 10.1093/molbev/msu400

49. Arellano-Saab, A., Bunsick, M., Al Galib, H., Zhao, W. D., Schuetz, S., Bradley, J. M., et al. (2021) Three mutations repurpose a plant karrikin receptor to a strigolactone receptor PROCEEDINGS OF THE NATIONAL ACADEMY OF SCIENCES OF THE UNITED STATES OF AMERICA 118, 10.1073/pnas.2103175118

50. Guercio, A. M., Palayam, M., and Shabek, N. (2023) Strigolactones: diversity, perception, and hydrolysis Phytochemistry Reviews 22, 339–359 10.1007/s11101-023-09853-4

51. de Saint Germain, A., Jacobs, A., Brun, G., Pouvreau, J. B., Braem, L., Cornu, D., et al. (2021) A Phelipanche ramosa KAI2 protein perceives strigolactones and isothiocyanates enzymatically Plant Communications 10.1016/j.xplc.2021.100166

52. Scaffidi, A., Waters, M. T., Sun, Y. K., Skelton, B. W., Dixon, K. W., Ghisalberti, E. L. et al. (2014) Strigolactone hormones and their stereoisomers signal through two related receptor proteins to induce different physiological responses in arabidopsis Plant Physiology 165, 1221–1232-1221–1232 10.1104/pp.114.240036

53. de Saint Germain, A., Clavé, G., Badet-Denisot, M. A., Pillot, J. P., Cornu, D., Le Caer, J. P., et al. (2016) An histidine covalent receptor and butenolide complex mediates strigolactone perception Nature Chemical Biology 12, 787–794-787–794 10.1038/nchembio.2147

54. Tsuchiya, Y., Yoshimura, M., Sato, Y., Kuwata, K., Toh, S., Holbrook-Smith, D., et al. (2015) Probing strigolactone receptors in Striga hermonthica with fluorescence Science 349, 10.1126/science.aab3831

55. Yao, J., Scaffidi, A., Meng, Y., Melville, K. T., Komatsu, A., Khosla, A. et al. (2021) Desmethyl butenolides are optimal ligands for karrikin receptor proteins New Phytologist 230, 1003–1016,

56. Yao, R., Ming, Z., Yan, L., Li, S., Wang, F., Ma, S. et al. (2016) DWARF14 is a non-canonical hormone receptor for strigolactone Nature 536, 469–473 10.1038/nature19073

57. Otwinowski, Z., and Minor, W. (1997) Processing of X-ray diffraction data collected in oscillation mode Methods in Enzymology 276, 307–326 10.1016/S0076-6879(97)76066-X

58. Lee, I. (2018) A missense allele of KARRIKIN-INSENSITIVE2 impairs ligand-binding and downstream signaling in Arabidopsis thaliana J Exp Bot 69, 10.1093/jxb/ery164

59. Adams, P. D. (2010) PHENIX: A comprehensive Python-based system for macromolecular structure solution Acta Crystallogr Sect D Biol Crystallogr 66, 10.1107/S0907444909052925

60. Emsley, P., Lohkamp, B., Scott, W. G., and Cowtan, K. (2010) Features and development of Coot Acta Crystallographica Section D: Biological Crystallography 66, 486–501 10.1107/S0907444910007493

61. Kumar, S., Stecher, G., Li, M., Knyaz, C., and Tamura, K. (2018) MEGA X: Molecular evolutionary genetics analysis across computing platforms Molecular Biology and Evolution 35, 1547–1549 10.1093/molbev/msy096

62. Larkin, M. A., Blackshields, G., Brown, N. P., Chenna, R., McGettigan, P. A., McWilliam, H. et al. (2007) Clustal W and Clustal X version 2.0 Bioinformatics 23, 2947–2948 10.1093/bioinformatics/btm404

63. Le, S. Q., and Gascuel, O. (2008) An improved general amino acid replacement matrix Mol Biol Evol 25, 1307–1320 10.1093/molbev/msn067

64. Weaver, S., Shank, S. D., Spielman, S. J., Li, M., Muse, S. V., and Kosakovsky Pond, S. L. (2018) Datamonkey 2.0: A Modern Web Application for Characterizing Selective and Other Evolutionary Processes Molecular Biology and Evolution 35, 773–777 10.1093/molbev/msx335

65. Eisenberg, D., Schwarz, E., Komaromy, M., and Wall, R. (1984) Analysis of membrane and surface protein sequences with the hydrophobic moment plot J Mol Biol 179, 125–142 10.1016/0022-2836(84)90309-7

