## Supporting information for "Structural insights into rice KAI2 receptor provide functional implications for perception and signal transduction"

**Figure S1.** Multiple sequence alignment of key KAI2/D14 sequences

**Figure S2.** Comparing overall structure and pocket residues of OsKAI2 with Arabidopsis and Striga

**Figure S3.** KAI2s used in this study and DSF data

**Figure S4.** KAI2s used in pulldown assays

**Table S1.** Data collection, phasing and refinement statistics

**Table S2.** Sequences used in amino acid tree and dN/dS analysis.

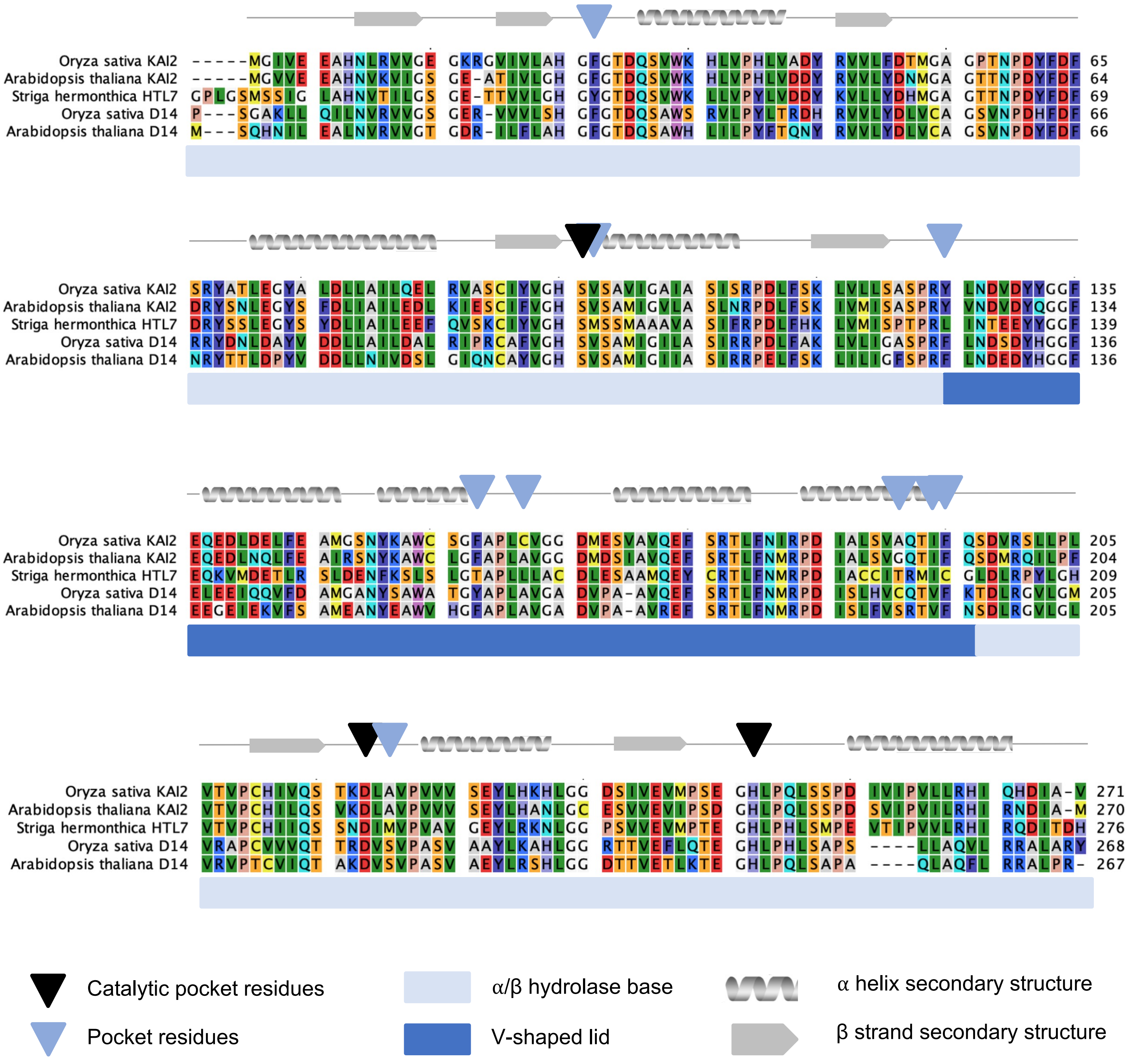

**Figure S1. Multiple sequence alignment of key KAI2/D14 sequences.** Amino acid alignment of four KAI2/D14s. Residues are colored in Rasmol standard colors. Numbers above residues refer to position in alignment. Black arrows refer to catalytic pocket residues S96, D218, and H247. Light blue arrows point out residues in pocket featured in main Fig. 1. OsKAI2 lid (light blue) and base (dark blue) domains are indicated below alignment. Secondary structure of OsKAI2 is shown above sequence in gray helices, beta strands, and non-secondary structure containing loops as lines.

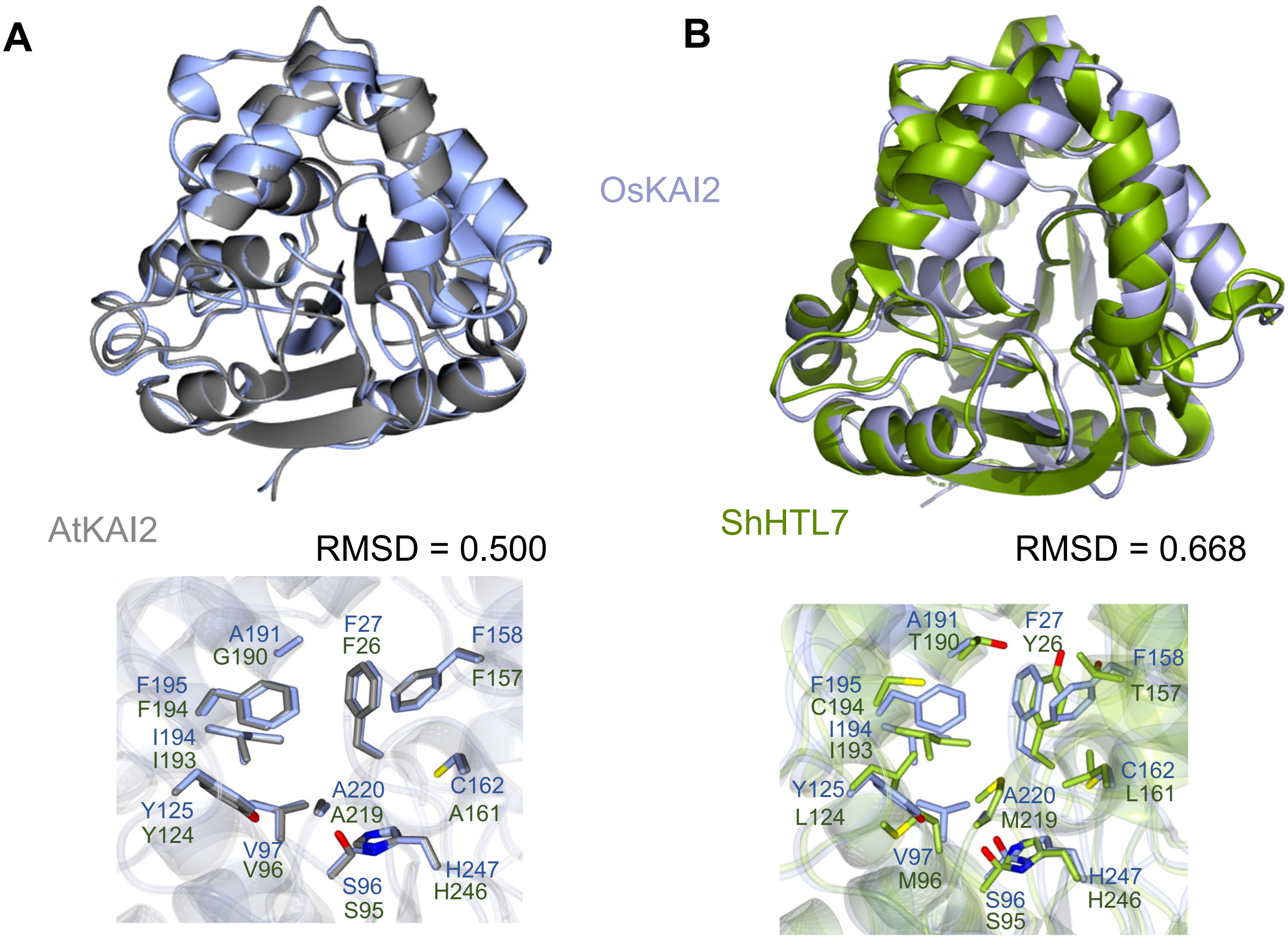

**C**

| | Surface area (A²) | Volume (A³) |
|---|---|---|
| OsKAI2 | 193.794 | 107.917 |
| AtKAI2 (4HRY) | 168.521 | 63.419 |
| ShD14 (5Z7Z) | 283.260 | 153.069 |
| OsD14 (4IH9) | 251.181 | 144.730 |
| ShHTL7 (5Z95) | 299.684 | 215.173 |
| PsKAI2(B) (7K2Z) | 234.425 | 125.377 |

**Figure S2. Comparing overall structure and pocket residues of OsKAI2 with Arabidopsis and Striga.** Superposition of OsKAI2 structure with **(A)** AtKAI2 and **(B)** ShHTL7 with corresponding RMSD for Cα. **(C)** Binding pocket dimensions are listed for a range of KAI2, D14, and HTL structures selected from the PDB as calculated by CASTp with a 1.4 Å radius probe.

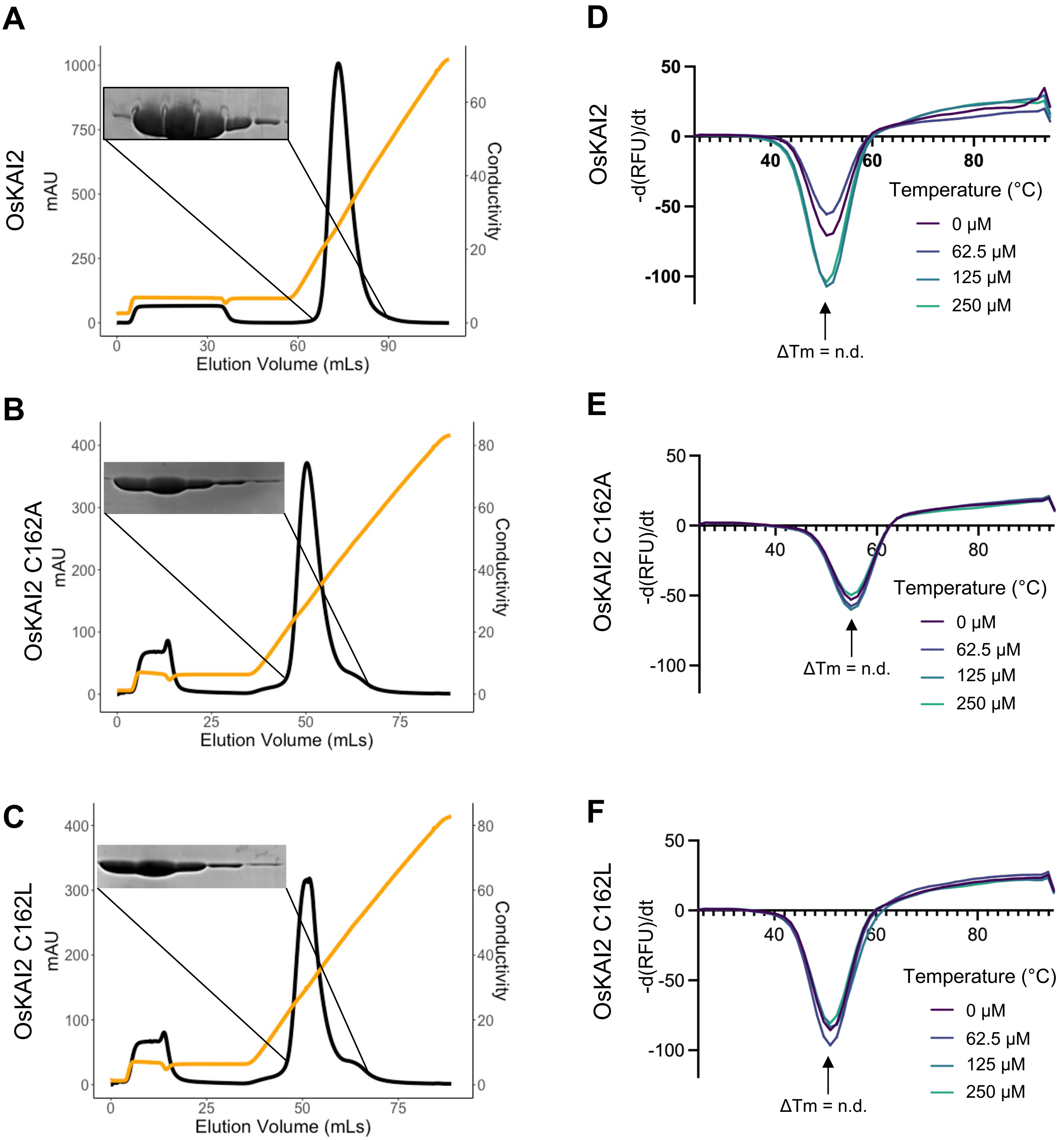

**Figure S3. KAI2s used in this study and DSF data. (A-C)** Purification anion exchange step for KAI2s used in study. **(D-F)** DSF analysis of OsKAI2, OsKAI2 C162A and OsKAI2 C162L in response to (+)-GR24.

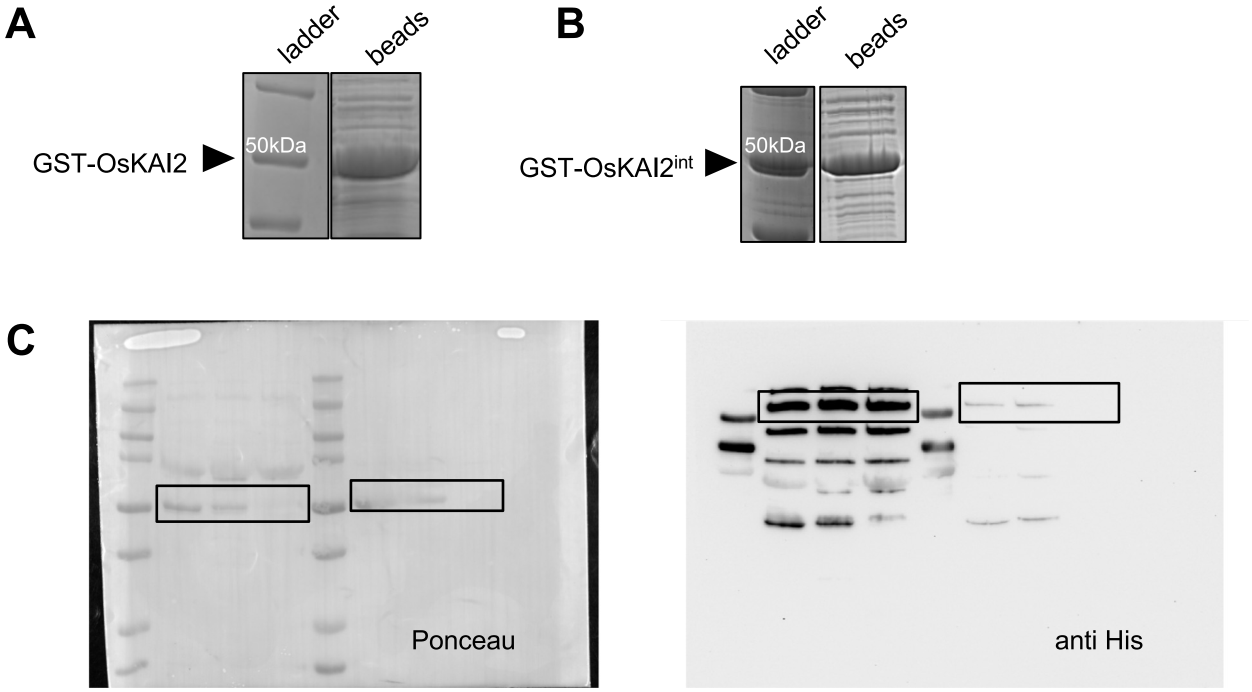

**Figure S4. KAI2s used in pulldown assays.** GST-KAI2 **(A)** or GST-KAI2$^{int}$ **(B)** -bound beads for use in pull-down assays. **(C)** Uncropped gel of pull-down ponceau (left) and anti His (right).

| | OsKAI2-'apo' | OsKAI2-MPD | AtD14 |
|---|---|---|---|
| **Data collection** | | | |
| Space group | $P1\ 2_1\ 1$ | $P1\ 2_1\ 1$ | $P1\ 2_1\ 1$ |
| Cell dimensions | | | |
| a, b, c (Å) | 37.64 85.39, 44.62 | 37.71, 86.48, 44.62 | 43.69, 69.27, 90.80 |
| α, β, γ (°) | 90.00, 114.91, 90.00 | 90.00, 114.94, 90.00 | 90.00 93.38 90.00 |
| Resolution (Å) | 42.73-1.68 (1.71-1.68) | 40.49-1.29 (1.31-1.29) | 45.32-2.00 (2.05-2.00) |
| $R_{sym}$ | 0.069 (0.422) | 0.051 (0.320) | 0.043 (0.317) |
| $\langle I / \sigma I \rangle$ | 16.0 (2.6) | 19.7 (2.6) | 14.6 (1.4) |
| Completeness (%) | 93.8 (84.1) | 99.3 (88.3) | 99.3 (91.7) |
| Redundancy | 3.5 (3.5) | 4.3 (3.2) | 5.8 (3.4) |
| **Refinement** | | | |
| Resolution (Å) | 1.68 | 1.29 | 2.00 |
| Unique reflections | 27040 (1274) | 64764 (2848) | 36532 (2459) |
| $R_{work}$ / $R_{free}$ (%) | 16.8/19.7 | 15.2/16.8 | 16.3/20.7 |
| No. atoms | 4158 | 4407 | 4357 |
| Protein | 4055 | 4138 | 4125 |
| Ligand/ion | - | 21 | - |
| Water | 103 | 245 | 232 |
| B-factors | | | |
| Protein | 22.7 | 14.7 | 33.85 |
| Ligand/ion | - | 15.1 | - |
| Water | 28.0 | 24.1 | 38.5 |
| R.m.s. deviations | | | |
| Bond lengths (Å) | 0.0084 | 0.0116 | 0.0067 |
| Bond angles (°) | 1.553 | 1.925 | 1.559 |
| Ramachandran favored (%) | 97.40 | 96.92 | 96.56 |
| Ramachandran allowed (%) | 2.60 | 3.08 | 3.06 |
| Ramachandran outliers (%) | 0 | 0 | 0.38 |
| PDB ID | 8VCZ | 8VD1 | 8VD3 |

**Table S1. Data collection, phasing and refinement statistics**. *Statistics for the highest-resolution shell are shown in parentheses.

| | | Species | Gene Name | Sequence ID | Source |
|---|---|---|---|---|---|
| MONOCOTs | Poaceae | Oryza sativa Japonica Group | probable esterase D14L | XP_015617518.1 | NCBI |
| MONOCOTs | Poaceae | Panicum hallii | probable esterase D14L | XP_025798102.1 | NCBI |
| MONOCOTs | Poaceae | Sorghum bicolor | probable esterase D14L | XP_002465088.1 | NCBI |
| MONOCOTs | Poaceae | Lolium rigidum | probable esterase D14L | XP_047090814.1 | NCBI |
| MONOCOTs | Poaceae | Brachypodium distachyon | probable esterase D14L | XP_003562367.1 | NCBI |
| MONOCOTs | Poaceae | Triticum dicoccoides | probable esterase D14L | XP_037430390.1 | NCBI |
| MONOCOTs | Poaceae | Aegilops tauschii subsp. strangulata | probable esterase D14L | XP_020178880.1 | NCBI |
| MONOCOTs | Poaceae | Triticum urartu | probable esterase D14L | XP_048570933.1 | NCBI |
| MONOCOTs | Poaceae | Hordeum vulgare subsp. vulgare | probable esterase D14L | XP_044981364.1 | NCBI |
| MONOCOTs | Poaceae | Setaria italica | probable esterase D14L | XP_004983951.1 | NCBI |
| MONOCOTs | Poaceae | Dichanthelium oligosanthes | putative esterase D14L | OEL18596.1 | NCBI |
| MONOCOTs | Poaceae | Oryza brachyantha | probable esterase D14L | XP_006650237.1 | NCBI |
| MONOCOTs | Poaceae | Panicum virgatum | probable esterase D14L | XP_039828974.1 | NCBI |
| MONOCOTs | Poaceae | Mascanthus sinensis | ESTERASE KAI2-RELATED | v7.1\|Misin01G312200.1.p | Phytozome |
| MONOCOTs | Poaceae | Mascanthus sinensis | ESTERASE KAI2-RELATED | v7.1\|Misin02G301000.1.p | Phytozome |
| MONOCOTs | Poaceae | Chasmanthium laxum | ESTERASE KAI2-RELATED | v1.1\|Chala.04G038000.1.p | Phytozome |
| MONOCOTs | Poaceae | Zea mays | ESTERASE KAI2-RELATED | RefGen_V4\|Zm00001d029590_P001 | Phytozome |
| MONOCOTs | Poaceae | Zea mays | ESTERASE KAI2-RELATED | RefGen_V4\|Zm00001d047335_P001 | Phytozome |
| MONOCOTs | Poaceae | Zea mays | ESTERASE KAI2-RELATED | RefGen_V4\|Zm00001d029590_P001 | Phytozome |
| MONOCOTs | Poaceae | Eleusine coracana | ESTERASE KAI2-RELATED | v1.1\|ELECO.r07.3AG0229120.1 | Phytozome |
| MONOCOTs | Poaceae | Eleusine coracana | ESTERASE KAI2-RELATED | v1.1\|ELECO.r07.3BG0274580.1 | Phytozome |
| MONOCOTs | Poaceae | Brachypodium sylvaticum | ESTERASE KAI2-RELATED | v2.1\|Brasyl.2G222900.1.p | Phytozome |
| MONOCOTs | Poaceae | Brachypodium arbuscula | ESTERASE KAI2-RELATED | v3.1\|Barbu.2G218500.1.p | Phytozome |
| MONOCOTs | Poaceae | Brachypodium hybridum | ESTERASE KAI2-RELATED | v2.1\|Bhyb26.D01G152100.1.p | Phytozome |
| MONOCOTs | Poaceae | Brachypodium hybridum | ESTERASE KAI2-RELATED | v2.1\|Bhyb26.S02G194500.1.p | Phytozome |
| MONOCOTs | Poaceae | Thinopyrum intermedium | ESTERASE KAI2-RELATED | v3.1\|Thint.J04G190800.1.p | Phytozome |
| MONOCOTs | Poaceae | Thinopyrum intermedium | ESTERASE KAI2-RELATED | v3.1\|Thint.S04G175500.1.p | Phytozome |
| MONOCOTs | Poaceae | Brachypodium mexicanum | ESTERASE KAI2-RELATED | v2.1\|Bmexi.02PG369700.1.p | Phytozome |
| MONOCOTs | Poaceae | Brachypodium mexicanum | ESTERASE KAI2-RELATED | v2.1\|Bmexi.02UG214100.1.p | Phytozome |
| MONOCOTs | Joinvilleaceae | Joinvillea ascendens | ESTERASE KAI2-RELATED | v1.1\|Joasc.01G055100.1.p | Phytozome |
| MONOCOTs | Joinvilleaceae | Joinvillea ascendens | ESTERASE KAI2-RELATED | v1.1\|Joasc.01G055000.1.p | Phytozome |
| MONOCOTs | Orchidaceae | Phalaenopsis equestris | probable esterase D14L | XP_020584281.1 | NCBI |
| MONOCOTs | Orchidaceae | Apostasia shenzhenica | putative esterase KAI2 | PKA58159.1 | NCBI |
| MONOCOTs | Orchidaceae | Dendrobium catenatum | probable esterase D14L | XP_020704619.1 | NCBI |
| MONOCOTs | Arecaceae | Elaeis guineensis | probable esterase D14L | XP_010905263.1 | NCBI |
| MONOCOTs | Arecaceae | Phoenix dactylifera | probable esterase D14L | XP_008787141.3 | NCBI |
| MONOCOTs | Cyperaceae | Carex littledalei | esterase D14L | KAF3323272.1 | NCBI |
| MONOCOTs | Musaceae | Musa acuminata subsp. malaccensis | probable esterase D14L | XP_009421091.1 | NCBI |
| MONOCOTs | Zingiberaceae | Zingiber officinale | probable esterase D14L | XP_042388420.1 | NCBI |
| MONOCOTs | Bromeliaceae | Ananas comosus | probable esterase D14L | XP_020082399.1 | NCBI |
| MONOCOTs | Dioscoreaceae | Dioscorea cayenensis subsp. rotundata] | probable esterase D14L | XP_039137355.1 | NCBI |
| MONOCOTs | Dioscoreaceae | Dioscorea alata | ESTERASE KAI2-RELATED | v2.1\|Dioal.13G030200.1.p | Phytozome |
| MONOCOTs | Asparagaceae | Asparagus officinalis | probable esterase D14L | XP_020242859.1 | NCBI |
| MONOCOTs | Asparagaceae | Agave tequilana var. Weber's Blue | ESTERASE KAI2-RELATED | v2.1\|AgateH1.08G198100.1.p | Phytozome |
| MONOCOTs | Acoraceae | Acorus americanus | ESTERASE KAI2-RELATED | v1.1\|Acora.05G132600.1.p | Phytozome |
| MONOCOTs | Araceae | Spirodela polyrhiza | ESTERASE KAI2-RELATED | S.polyrhiza v2\|Spipo2G0095700 | Phytozome |
| BASAL ANGIOSPERMS and OTHER | Nymphaeaceae | Nymphaea colorata | probable esterase D14L | XP_031494429.1 | NCBI |
| BASAL ANGIOSPERMS and OTHER | Amborellaceae | Amborella trichopoda | probable esterase D14L | XP_006829226.1 | NCBI |
| BASAL ANGIOSPERMS and OTHER | Lauraceae | Cinnamomum micranthum f. kanehirae | putative esterase D14L | RWR81441.1 | NCBI |
| BASAL ANGIOSPERMS and OTHER | Magnoliaceae | Liriodendron tulipifera | ESTERASE KAI2-RELATED | v1.1\|Litul.17G055800.1.p | Phytozome |
| EUDICOTS | Proteaceae | Telopea speciosissima | probable esterase KAI2 | XP_043715070.1 | NCBI |
| EUDICOTS | Proteaceae | Macadamia integrifolia | probable esterase KAI2 | XP_042493687.1 | NCBI |
| EUDICOTS | Malvaceae | Gossypium hirsutum | probable esterase D14L | XP_016719056.1 | NCBI |
| EUDICOTS | Malvaceae | Hibiscus syriacus | probable esterase KAI2 | XP_038996839.1 | NCBI |
| EUDICOTS | Malvaceae | Gossypium australe | probable esterase D14L | KAA3457435.1 | NCBI |
| EUDICOTS | Malvaceae | Theobroma cacao | probable esterase D14L | XP_017969914.1 | NCBI |
| EUDICOTS | Malvaceae | Herrania umbratica | probable esterase KAI2 | XP_021274049.1 | NCBI |
| EUDICOTS | Actinidiaceae | Actinidia eriantha | probable esterase D14L | XP_057514426.1 | NCBI |
| EUDICOTS | Amaranthaceae | Chenopodium quinoa | probable esterase KAI2 | XP_021714410.1 | NCBI |
| EUDICOTS | Amaranthaceae | Beta vulgaris subsp. Vulgaris | probable esterase D14L | XP_010679577.1 | NCBI |
| EUDICOTS | Amaranthaceae | Spinacia oleracea | ESTERASE KAI2-RELATED | S.oleracea Spov3\|Spov3_chr3.01003 | Phytozome |
| EUDICOTS | Amaranthaceae | Amaranthus hypochondriacus | ESTERASE KAI2-RELATED | v2.1\|AH006186-RA | Phytozome |
| EUDICOTS | Celastraceae | Tripterygium wilfordii | probable esterase KAI2 | XP_038723957.1 | NCBI |
| EUDICOTS | Rhamnaceae | Ziziphus jujuba var. spinosa | probable esterase KAI2 | XP_048335111.1 | NCBI |
| EUDICOTS | Theaceae | Camellia lanceoleosa | probable esterase D14L | KAI8028962.1 | NCBI |
| EUDICOTS | Myricaceae | Morella rubra | probable esterase D14L | KAB1214871.1 | NCBI |
| EUDICOTS | Anacardiaceae | Pistacia vera | probable esterase KAI2 | XP_031277949.1 | NCBI |
| EUDICOTS | Caricaceae | Carica papaya | probable esterase KAI2 | XP_021908336.1 | NCBI |
| EUDICOTS | Papaveraceae | Papaver somniferum | probable esterase KAI2 | XP_026394197.1 | NCBI |
| EUDICOTS | Solanaceae | Solanum lycopersicum | ESTERASE KAI2-RELATED | ITAG5.0\|Solyc02T002798.2 | Phytozome |
| EUDICOTS | Solanaceae | Solanum lycopersicum | ESTERASE KAI2-RELATED | ITAG5.0\|Solyc02T000856.1 | Phytozome |
| EUDICOTS | Solanaceae | Solanum tuberosum | ESTERASE KAI2-RELATED | v6.1\|Soltu.DM.02G007700.1 | Phytozome |
| EUDICOTS | Solanaceae | Solanum tuberosum | ESTERASE KAI2-RELATED | v6.1\|Soltu.DM.02G007690.1 | Phytozome |
| EUDICOTS | Solanaceae | Solanum tuberosum | ESTERASE KAI2-RELATED | v6.1\|Soltu.DM.02G028010.1 | Phytozome |
| EUDICOTS | Brassicaceae | Arabidopsis thaliana | KAI2 | AT4G37470.1 | TAIR |
| EUDICOTS | Brassicaceae | Capsella rubella | ESTERASE KAI2-RELATED | v1.1\|Carub.0007s0281.1.p | Phytozome |

| | | Species | Gene Name | Sequence ID | Source |
|---|---|---|---|---|---|
| EUDICOTS | Phrymaceae | Erythranthe guttata | KAI2a | XP_012856373.1 | NCBI |
| EUDICOTS | Phrymaceae | Erythranthe guttata | KAI2b | XP_012853813.1 | NCBI |
| EUDICOTS | Phrymaceae | Erythranthe guttata | KAI2c | XP_012840794.1 | NCBI |
| EUDICOTS | Vitaceae | Vitis vinifera | probable esterase KAI2 | XP_002284043.1 | NCBI |
| EUDICOTS | Salicaceae | Populus trichocarpa | KAI2a | XP_002307293.1 | NCBI |
| EUDICOTS | Salicaceae | Populus trichocarpa | KAI2b | XP_006380654.1 | NCBI |
| EUDICOTS | Rosaceae | Prunus persica | probable esterase KAI2 | XP_007205717.1 | NCBI |
| EUDICOTS | Fabaceae | Medicago truncatula | KAi2a | Mt4.0v1\|Medtr4g095310.1 | Phytozome |
| EUDICOTS | Fabaceae | Medicago truncatula | KAI2b | Mt4.0v1\|Medtr5g016150.1 | Phytozome |
| EUDICOTS | Fabaceae | Lotus Japonicus | KAI2a | Lj1.0v1\|Lj2g0007403.1 | Phytozome |
| EUDICOTS | Fabaceae | Lotus Japonicus | KAI2b | Lj1.0v1\|Lj4g0019758.1 | Phytozome |
| EUDICOTS | Fabaceae | Cicer arietinum | ESTERASE KAI2-RELATED | v1.0\|Ca_02196 | Phytozome |
| EUDICOTS | Fabaceae | Cicer arietinum | ESTERASE KAI2-RELATED | v1.0\|Ca_09326 | Phytozome |
| EUDICOTS | Fabaceae | Glycine max | ESTERASE KAI2-RELATED | v2.1\|GmISU01.17G149900.1.p | Phytozome |
| EUDICOTS | Fabaceae | Glycine max | ESTERASE KAI2-RELATED | v2.1\|GmISU01.17G150000.1.p | Phytozome |
| EUDICOTS | Fabaceae | Glycine max | ESTERASE KAI2-RELATED | v2.1\|GmISU01.01G152200.1.p | Phytozome |
| EUDICOTS | Fabaceae | Glycine max | ESTERASE KAI2-RELATED | v2.1\|GmISU01.11G045700.1.p | Phytozome |
| EUDICOTS | Fabaceae | Glycine max | ESTERASE KAI2-RELATED | v2.1\|GmISU01.05G083300.1.p | Phytozome |
| EUDICOTS | Fabaceae | Pisum sativum | KAI2a | | Guercio et al 2022 |
| EUDICOTS | Fabaceae | Pisum sativum | KAI2b | | Guercio et al 2022 |
| EUDICOTS | Crassulaceae | Kalanchoe fedtschenkoi | ESTERASE KAI2-RELATED | v1.1\|Kaladp0034s0001.1.p | Phytozome |
| EUDICOTS | Portulacaceae | Portulaca amilis | ESTERASE KAI2-RELATED | v1.0\|FUN_030516-T1 | Phytozome |
| EUDICOTS | Crassulaceae | Kalanchoe laxiflora | ESTERASE KAI2-RELATED | v3.1\|Kalaxd.05G066100.1.p | Phytozome |
| EUDICOTS | Fagaceae | Castanea dentata | ESTERASE KAI2-RELATED | v1.1\|Caden.02G181900.1.p | Phytozome |
| EUDICOTS | Ranunculaceae | Aquilegia coerulea | ESTERASE KAI2-RELATED | v3.1\|Aqcoe6G293500.1.p | Phytozome |
| EUDICOTS | Ranunculaceae | Aquilegia coerulea | ESTERASE KAI2-RELATED | v3.1\|Aqcoe6G293700.1.p | Phytozome |
| EUDICOTS | Ranunculaceae | Aquilegia coerulea | ESTERASE KAI2-RELATED | v3.1\|Aqcoe6G293600.1.p | Phytozome |
| EUDICOTS | Ranunculaceae | Aquilegia coerulea | ESTERASE KAI2-RELATED | v3.1\|Aqcoe1G368100.1.p | Phytozome |
| EUDICOTS | Hydrangeaceae | Hydrangea quercifolia | ESTERASE KAI2-RELATED | v1.1\|Hyque.05G069100.1.p | Phytozome |
| EUDICOTS | Hydrangeaceae | Hydrangea quercifolia | ESTERASE KAI2-RELATED | v1.1\|Hyque.05G069300.1.p | Phytozome |
| EUDICOTS | Asteraceae | Helianthus annuus | ESTERASE KAI2-RELATED | r1.2\|HanXRQChr12g0373271 | Phytozome |
| EUDICOTS | Asteraceae | Helianthus annuus | ESTERASE KAI2-RELATED | r1.2\|HanXRQChr11g0322481 | Phytozome |
| EUDICOTS | Asteraceae | Lactuca sativa | ESTERASE KAI2-RELATED | V8\|Lsat_1_v5_gn_4_162120.1 | Phytozome |
| EUDICOTS | Asteraceae | Lactuca sativa | ESTERASE KAI2-RELATED | V8\|Lsat_1_v5_gn_4_162080.1 | Phytozome |
| EUDICOTS | Oleaceae | Olea europaea | ESTERASE KAI2-RELATED | v1.0\|Oeu011610.1 | Phytozome |
| EUDICOTS | Oleaceae | Olea europaea | ESTERASE KAI2-RELATED | v1.0\|Oeu040607.1 | Phytozome |
| EUDICOTS | Apiaceae | Daucus carota | ESTERASE KAI2-RELATED | v2.0\|DCAR_010837 | Phytozome |
| EUDICOTS | Apiaceae | Daucus carota | ESTERASE KAI2-RELATED | v2.0\|DCAR_017602 | Phytozome |
| EUDICOTS | Rutaceae | Citrus sinensis | ESTERASE KAI2-RELATED | v1.1\|orange1.1g046596m | Phytozome |
| EUDICOTS | Betulaceae | Betula platyphylla | ESTERASE KAI2-RELATED | v1.1\|BPChr06G22594 | Phytozome |
| EUDICOTS | Betulaceae | Betula platyphylla | ESTERASE KAI2-RELATED | v1.1\|BPChr06G22550 | Phytozome |
| EUDICOTS | Betulaceae | Betula platyphylla | ESTERASE KAI2-RELATED | B.platyphylla v1.1\|BPChr06G22593 | Phytozome |
| EUDICOTS | Cucurbitaceae | Cucumis sativus | ESTERASE KAI2-RELATED | v1.0\|Cucsa.102900.1 | Phytozome |
| EUDICOTS | Cucurbitaceae | Cucumis sativus | ESTERASE KAI2-RELATED | v1.0\|Cucsa.367130.1 | Phytozome |
| EUDICOTS | Orobanchaceae | Lindenbergia philippensis | ESTERASE KAI2-RELATED | v1.1\|Liphi.09G063400.1.p | Phytozome |
| EUDICOTS | Orobanchaceae | Lindenbergia philippensis | ESTERASE KAI2-RELATED | v1.1\|Liphi.04G117700.1.p | Phytozome |
| EUDICOTS | Orobanchaceae | Striga hermonthica | HTL1 | | Toh et al 2015 |
| EUDICOTS | Orobanchaceae | Striga hermonthica | HTL2 | | Toh et al 2015 |
| EUDICOTS | Orobanchaceae | Striga hermonthica | HTL3 | | Toh et al 2015 |
| EUDICOTS | Orobanchaceae | Striga hermonthica | HTL4 | | Toh et al 2015 |
| EUDICOTS | Orobanchaceae | Striga hermonthica | HTL5 | | Toh et al 2015 |
| EUDICOTS | Orobanchaceae | Striga hermonthica | HTL6 | | Toh et al 2015 |
| EUDICOTS | Orobanchaceae | Striga hermonthica | HTL7 | | Toh et al 2015 |
| EUDICOTS | Orobanchaceae | Striga hermonthica | HTL8 | | Toh et al 2015 |
| EUDICOTS | Orobanchaceae | Striga hermonthica | HTL9 | | Toh et al 2015 |
| EUDICOTS | Orobanchaceae | Striga hermonthica | HTL10 | | Toh et al 2015 |
| EUDICOTS | Orobanchaceae | Striga hermonthica | HTL11 | | Toh et al 2015 |
| EUDICOTS | D14s | Arabidopsis thaliana | D14 | AT3G03990.1 | TAIR |

**Table S2. All sequences used in amino acid tree and dN/dS analysis.**
